# ABF2 and bZIP2 remodel root system architecture under combined phosphate deficiency and salinity in *Arabidopsis*

**DOI:** 10.64898/2026.08.28.747924

**Authors:** H. Salinas-Grenet, N.R. Johnson, M.A. Ibeas, T.C. Moyano, J. Pérez-Díaz, R. Achá-Escobar, L. Ahumada-Langer, AR Kim, V. Berdion Gabarain, G. Vásquez-Marambio, N Perrimon, J.M. Álvarez, J.M. Estevez

## Abstract

Climate change is degrading arable soils and imposing chronic abiotic stress on plants. As the primary interface with the soil, the root system is especially vulnerable, and plants respond by remodelling their root system architecture (RSA). Most studies have examined single stresses in isolation and so fail to capture the multifactorial environments in which plants actually grow. Rising salinity impairs development by compromising cellular integrity and triggering cytotoxic responses, and it simultaneously restricts uptake of phosphorus, a macronutrient required for nucleic acid and protein synthesis. Combined stresses can amplify these effects, yet how plants adjust RSA to cope with them remains poorly understood. Here we performed a meta-analysis of public transcriptomic datasets for salinity and phosphate deficiency in Arabidopsis thaliana. Integrating these data with protein-protein interaction modelling and DNA-binding (DAP-seq) analyses, we identified two basic leucine zipper transcription factors, ABF2 and bZIP2, as candidate integrators of the combined-stress response. Characterization of *abf2*, *bzip2* and double mutants revealed RSA alterations and allowed us to identify downstream targets of each factor. Our findings offer new insight into root adaptation under combined stress and represent one of the few comprehensive analyses of how multiple abiotic stressors jointly shape root development and plant fitness.

**Highlight:** Data mining of public transcriptomes, network inference and AlphaFold-Multimer screening identify ABF2 and bZIP2 as interacting regulators of root remodelling under combined stress.

## Introduction

Abiotic stress can substantially impair plant function and fitness by forcing physiological readjustment. Nutrient availability, which is largely set by soil quality, drives molecular-level changes by reshaping metabolic pathways, cellular sugar content, and source–sink dynamics, among other processes (Laudicina et al., 2023). These changes have direct consequences for commercial production, for instance by degrading fruit quality and lowering soluble-solid content and, with it, organoleptic value (Dhanyasree et al., 2022). Understanding the increasingly challenging scenarios imposed by climate change is therefore essential, particularly in the context of nutrient deficiency and soil erosion, which continuously expose plants to multiple, co-occurring abiotic stresses (Oishy et al., 2025).

The root is the plant organ most directly affected by abiotic stress, acting at once as a barrier to and a sensor of soil-derived signals. Root growth and development are tightly regulated and continuously remodeled; roots are central to nutrient uptake and contribute substantially to stress mitigation within the soil. To meet these challenges, roots can modify their spatial distribution and morphology, a property captured by the concept of root system architecture (RSA) (Damena, 2023; Karlova et al., 2021). RSA is defined as the spatial configuration of the root system and the geometry of its individual axes (Lynch, 1995) and comprises the primary root (PR), lateral roots (LR), and root hairs (RH) (Jung and McCouch, 2013; Liu, 2021). These components are highly responsive to environmental cues, and RSA remodeling can enhance or reshape fitness through interactions with the microbiome, improved nutrient acquisition, and optimized water uptake (Lynch, 1995; Van Zelm et al., 2020).

RSA is strongly affected by saline stress. In the PR, salinity disrupts the apical meristem responsible for cell division and elongation (Farooq et al., 2024) and promotes salt-avoidance growth through redistribution of auxin gradients mediated by PIN-family auxin efflux carriers (Zwiewka et al., 2015). Salt stress also constrains LR emergence via ionic-stress signaling through the Salt Overly Sensitive (SOS) pathway, in which *SOS3* enhances auxin biosynthesis and transport in response to salinity, yielding shorter, less branched LRs (Zhao et al., 2011). LR density is likewise shaped by nutrient status: low phosphate availability in in vitro systems promotes lateral branching and higher LR density, whereas nitrate deficiency has the opposite effect (Julkowska et al., 2014; Zhao et al., 2011; Zolla et al., 2010), pointing to extensive crosstalk between stress responses.

Salinity directly impairs RH development by suppressing elongation and reducing density, consequences of disrupted ionic homeostasis, altered Ca2+ fluxes, and loss of the epidermal cells that give rise to RHs (Dinneny et al., 2008; Wang et al., 2008). ABA-responsive element binding factors (ABFs) act upstream of ROOT HAIR DEFECTIVE 6 (RHD6) in this response: ABF3 interacts physically with RHD6 and suppresses its activation of ROOT HAIR DEFECTIVE SIX-LIKE 4 (RSL4) under saline conditions (Jin et al., 2023). *RSL4* acts as a master regulator of root hair formation, controlling both RH initiation and growth (Yi et al., 2010). High salinity further perturbs phosphate (Pi) homeostasis and aggravates Pi-uptake limitations in species such as melon and lupin (Navarro et al., 2001; Treeby and van Steveninck, 1988). Salinity thus directly compromises Pi acquisition, generating a combined-stress scenario that remains largely unexplored.

Phosphate is an essential, finite, and non-renewable macronutrient central to photosynthesis, respiration, nucleic acid biosynthesis, and protein synthesis (Rubio et al., 2009; Noe et al., 2020). Soil Pi deficiency, with available concentrations below 1–10 µM, affects more than 40% of arable land worldwide and necessitates widespread fertilizer use. Yet only a small fraction of applied Pi (10–20%) is taken up by crops; the remainder becomes immobilized in soil or is lost to the environment through runoff (Lambers et al., 2006; Kong et al., 2018).

Root-mediated Pi acquisition relies on the secretion of exudates—including acid phosphatases, RNases, carboxylates, and protons—that solubilize immobilized Pi for uptake (Neumann and Römheld, 1999; Ryan et al., 2001; Hurley et al., 2010; Wang et al., 2011; Tian and Liao, 2015; Wang and Liu, 2018; Wang and Lambers, 2020). Pi deficiency strongly reshapes RSA and, in turn, fitness by altering shoot-to-root ratios and reducing yield in major crops (Rubio et al., 2009; Vance et al., 2003). To cope with Pi starvation, plants deploy several adaptive strategies, prominent among them the optimization of RSA.

The root tip serves as a Pi-availability sensor: under low Pi, cell division declines and quiescent-center identity is lost (Huang et al., 2020). The root apical meristem (RAM) tunes the magnitude of the Pi response through genes such as *LOW PHOSPHATE ROOT 1 (LPR1)* and the *ATPase PHOSPHATE DEFICIENCY RESPONSE 2 (PDR2)*, both identified by Quantitative Trait Locus analysis (Ticconi et al., 2004, Ticconi et al., 2009). Under Pi limitation, PDR2 regulates SCARECROW (SCR), a transcription factor that governs cell patterning and elongation and ultimately PR length (Reymond et al., 2006). Local Pi sensing at the meristem is further modulated by iron-dependent callose deposition (Müller et al., 2015). Mutants defective in *PDR2* and other Pi-sensing genes show altered LR development, with increased LR number and length that favor nutrient capture (Péret et al., 2011). Pi deficiency also markedly stimulates RH development, increasing both density and length. Morphologically, this root hair response relies on the strong induction of downstream effectors like the master regulator *RSL4*. In parallel, the broader systemic transcriptomic response is governed by central regulators such as *PHOSPHATE STARVATION RESPONSE 1 (PHR1)* and *PHR1-LIKE 1 (PHL1)*, which are essential for intracellular Pi homeostasis (Bustos et al., 2010). Accordingly, phr1/phl1 double mutants fail to mount a proper Pi-starvation transcriptional response (Bustos et al., 2010). Collectively, phosphorus is fundamental to plant physiology and to resistance against abiotic stress.

Plants sense environmental stress and modulate Pi signaling to tolerate stressors such as drought, heat, heavy-metal toxicity, and salinity (Khan et al., 2023). These pathways are therefore deeply interconnected, yet information on their combined effects remains scarce. Ibeas et al. (2024) recently provided the first comprehensive treatment of this question by integrating a meta-analysis of single-cell (scRNA-seq) and bulk RNA-seq data from root hairs exposed to abscisic acid, low phosphate, and high salinity. That work identified a set of genes that converge under phosphate and salt stress and are linked to root hair development.

In the present study, we applied a transcriptomic meta-analysis and data-mining strategy to identify and prioritize novel transcription factors involved in RSA remodeling under combined low-phosphate and salt stress. Through gene regulatory network reconstruction and AlphaFold-Multimer protein–protein interaction analyses, we identified two basic leucine zipper (bZIP) transcription factors, ABF2 and bZIP2, as strong candidates that physically interact to coordinate this response. We subsequently characterized the *abf2*, *bzip2*, and double mutants, revealing an additive or synergistic requirement for these factors to sustain lateral root emergence and root hair elongation under combined stress. Finally, by integrating DNA Affinity Purification sequencing (DAP-seq) with RNA-seq, we mapped their direct downstream targets, elucidating a three-module regulatory mechanism that links ABA-mediated dimerization to the specific regulation of root architecture, osmoprotection, and nutrient transport. Together, our results provide the first mechanistic evidence for how these novel TF complexes govern RSA remodeling under the complex constraints of combined low-phosphate and salt stress.

## Materials and Methods

### Data mining of relevant studies

Sequencing libraries relevant to phosphate and salt treatments in Arabidopsis thaliana were identified through a targeted data mining approach. Searches were performed with the NCBI EDirect tools using the broad query “(Phosphate OR Salt) AND Arabidopsis [orgn:txid3702]” against the BioProject, SRA, GDS, and BioSample databases, which were then merged into a single library dataset. Retrieved libraries were systematically categorized and annotated with metadata, including tissue type, developmental stage at harvest, treatment duration, and methodological details of the applied stress. Only datasets linked to a published article or preprint, or those with sufficiently detailed metadata, were retained. To ensure robustness, we kept only libraries from treatments (time-course or presence/absence) in A. thaliana Col- 0 wild type, excluding those limited to mutant-versus-wild-type comparisons, and required a minimum of two replicates per condition. Finally, libraries were filtered to retain only those addressing phosphate-related stress (deficiency, starvation, or low supply) or salt stress in any form of salt addition.

### Meta-RNA-seq analysis pipeline

Filtered libraries were downloaded with fasterq-dump 3.2.1 (NCBI SRA Toolkit), and read counts were verified against the associated metadata. Adapter trimming used cutadapt 5.2 (Martin, 2011) with the command cutadapt -q 25,25 -m 30 against a set of known RNA-seq adapters. Gene quantification was performed with kallisto quant v0.46.1 (Bray et al., 2016) against the ARAPORT11 transcriptome, using -b 100, -l 50, and -s 20 for single-end libraries. Differential-expression comparisons (“experiments”) were defined from the control-versus-treatment conditions above, prioritizing presence/absence designs and, where unavailable, time-course data referenced to time zero. Differential expression was assessed with DESeq2 (Love et al., 2014), and DE genes were called by Wald test (FDR < 0.05, no fold-change threshold). DE gene lists were then integrated across experiments to identify shared DEGs (sDEGs) Integration used a Monte Carlo simulation that tested the likelihood of a gene being DE across multiple experiments (Johnson et al., 2026) setting the threshold at the level exceeded by fewer than 10% of randomly selected gene sets of comparable size yielded cutoffs of five experiments for phosphate and six for salt. To further limit study-specific bias, genes were additionally required to be DE in at least two independent studies. Genes were then classified per treatment as up-regulated, down-regulated, bidirectionally regulated, or not an sDEG.

### Genesect analyses

We performed set comparisons with the genesectr package (github.com/nateyjay/genesectr), following methodologies established in VirtualPlant (Katari et al., 2010) and ConnecTF (Brooks et al., 2021). This tool applies Fisher’s exact test to evaluate enrichment or depletion between gene sets pairwise. All analyses were one-sided, and p-values were corrected with the Benjamini–Hochberg procedure.

### Network analysis

Gene regulatory networks (GRNs) were constructed independently for each treatment with the GENIE3 algorithm (Huynh-Thu et al., 2010). Inputs were raw counts from filtered libraries grouped by treatment, with separate networks for control and treatment conditions. The TF reference list was obtained from PlantTFDB v5.0 (Tian et al., 2020). GENIE3 outputs were filtered to retain the top 10% of interactions per target gene based on interaction weight, and the control and treatment networks were then merged by averaging interaction weights. To increase confidence, we retained only interactions supported by DAP-seq evidence from the Plant Cistrome Database (O’Malley et al., 2016), yielding treatment-specific, high-confidence GRNs.

### Identification of putative dimeric interactions between transcription factors

To identify putative dimeric interactions between transcription factors (TFs), genomic coordinates of TF-binding sites were obtained from the DAP-seq dataset reported by O’Malley et al. (2016). Binding sites corresponding to the TFs of interest were retrieved in BED format and intersected with promoter regions defined upstream of the transcription start site (TSS) of the analyzed genes. For each promoter, binding sites associated with individual TFs were identified and the genomic distances between sites corresponding to different TFs were calculated. Pairs of TF-binding sites occurring within the same promoter and separated by less than the predefined distance threshold were considered candidate spatially proximal binding events potentially compatible with dimeric or cooperative TF interactions. Similarly, closely spaced binding sites assigned to the same TF were considered as potential configurations compatible with homodimeric binding. Because the underlying DAP-seq experiments were performed independently for each TF, these associations were interpreted as putative dimeric interactions inferred from binding-site proximity rather than as direct evidence of protein–protein interaction.

### Plant material and growth conditions

All Arabidopsis thaliana lines were in the Columbia-0 (Col-0) background. For *abf2*, we used the SALK_002984 mutant line (Kim et al., 2004). For *hb6*, two insertional mutant lines were used: SALK_123279 and SALK_055797. For *bzip2*, the SAIL_619_F11.1 insertional line was used. To genotype the SALK lines, the LB1.3 T-DNA border primer was used to detect the mutant allele, while primers spanning the insertion site were used to amplify the wild-type (WT) allele. A similar strategy was employed for the SAIL bzip2 line, using the LB3 primer for the mutant allele. To generate the *abf2/bzip2* double mutant, the *abf2* and *bzip2* single mutants described above were crossed, and homozygous double mutants were selected via PCR genotyping using the respective insertion primers.

Seeds were surface-sterilized and stratified in the dark at 4 °C for 48 h before germination on 0.5× MS- MES (Duchefa, Haarlem, The Netherlands) supplemented with 0.8% Plant Agar™ (Duchefa) in 120 × 120 mm square Petri dishes (Deltalab, Barcelona, Spain), under continuous light (120 µmol m⁻² s⁻¹). For RSA phenotyping, seeds were stratified at 4 °C for 48 h and grown on half-strength (0.5×) MS agar at 22 °C under continuous light (120 µmol m⁻² s⁻¹) for 7 days. Seven-day-old seedlings were then transferred for 3 days to one of the following treatments: basal medium (0.5× MS), salt stress (100 mM NaCl), low phosphate (5 µM, supplied as K₂HPO₄), or combined stress (100 mM NaCl + 5 µM Pi), after which RSA parameters were measured. The pH of all media was adjusted to approximately 5.8 using 1 M KOH.

### Measurement of RSA traits under combined stress

RSA traits comprised primary root length (cm), total lateral root length (cm), number of lateral roots, lateral root density (calculated per seedling as lateral root number divided by primary root length prior to averaging), and root hair length. Measurements were made in ImageJ (Schneider et al., 2012). For macroscopic seedling traits, ten seedlings (n = 10) were analyzed per biological replicate across three replicates (total n = 30 seedlings per genotype and condition). For root hair length, the 10 longest root hairs were measured per seedling, yielding a total of 300 root hairs per genotype and condition. Whole-plant images were captured on an EPSON Perfection V8 scanner, and root hair images on a Leica EZ4 HD stereomicroscope with LAS EZ software.

### Protein-complex prediction with AlphaFold-Multimer

Protein-complex prediction used LocalColabFold v1.5.2 (Mirdita et al., 2022), which integrates AlphaFold-Multimer v2.3.1 (Evans et al., 2022) with MMseqs2 for multiple-sequence-alignment generation. Computations were run on the Harvard O2 high-performance computing cluster. For each complex, five models were generated with five recycling iterations each. In total, 120 pairwise combinations were screened, representing 105 heterotypic pairs and 15 homodimers. The integrated Local Interaction Score (iLIS) and integrated Local Interaction Area (iLIA) were computed to evaluate candidate interactions (Kim et al., 2026). A baseline positive interaction was defined using a reference-set-derived empirical threshold of iLIS ≥ 0.223. To further filter for the most robust assemblies, a more stringent cut-off of iLIS ≥ 0.551 was applied (Kim et al., 2026); the code is available at github.com/flyark/AFM-LIS. Predicted structures were visualized in ChimeraX (Meng et al., 2023). The TFs analyzed were WRKY33 (AT2G38470, Q8S8P5), RAV1 (AT1G13260, Q9ZWM9), HHO3 (AT1G25550, Q9FPE8), AT1G19000 (Q9LMC7), MYB13 (AT1G06180, Q9LNC9), HB6 (AT2G22430, P46668), STZ (AT1G27730, Q96289), DREB (AT1G01250, Q1ECI2), ERF/DREB subfamily A-6 (AT1G36060, Q9SKW5), ABF2 (AT1G45249, Q9M7Q4), bZIP21 (AT1G08320, Q93XM6), bZIP2 (AT2G18160, Q9SI15), AT1G74840 (A0A178WGE9), RAP2.12/ERF74 (AT1G53910, Q9SSA8), and WRKY40 (AT1G80840, Q9SAH7).

### Promoter motif-spacing analysis

To identify putative dimeric interactions between transcription factors, genomic coordinates of TF-binding sites for ABF2, bZIP2, and HB6 were obtained in BED format from the DAP-seq dataset (O’Malley et al., 2016). These binding sites were intersected with promoter regions, defined as the [insert exact window, e.g., 1000 bp] region upstream of the annotated transcription start site (TSS) of each target gene in ARAPORT11. For each promoter, the genomic distances between binding sites were calculated. Pairs of TF-binding sites separated by less than a predefined distance threshold [insert exact threshold, e.g., 3-10 bp] were considered candidate spatially proximal binding events compatible with cooperative TF interactions. Targets carrying two closely spaced binding sites assigned to the same TF were considered as potential configurations compatible with homodimeric binding, whereas targets carrying sites for different TFs (e.g., ABF2 and bZIP2) within the same spacing window were assigned to a heterodimeric configuration. Because the underlying DAP-seq experiments were performed independently for each TF, these associations were interpreted as putative dimeric interactions inferred from binding-site proximity rather than as direct evidence of protein–protein interaction.

## Statistical analysis

All RSA traits were analysed in GraphPad Prism v8.0.1. Data were assessed for normality (Shapiro-Wilk test) and for homogeneity of variances before parametric testing. Differences between genotypes and treatments were tested by two-way ANOVA (genotype × treatment) followed by Tukey’s multiple-comparison test; all tests were two-sided and P values are adjusted for multiple comparisons. Measurements were taken from distinct seedlings (biological replicates); no technical replicates were averaged. For macroscopic traits n = 30 seedlings per genotype and condition (10 seedlings × 3 independent biological replicates); for root hair length n = 300 root hairs per genotype and condition (the 10 longest root hairs of each of 30 seedlings). Because root hairs measured on the same seedling are not independent, this nesting is stated explicitly here and in the figure legends. For the summarized phenotypic heatmap (Fig. 5C) the raw data were normalized as the log2 fold-change relative to the contemporaneous Col-0 control and visualized in Python v3.9 with seaborn v0.11 and SciPy.

## Results

### An integrated strategy for identifying novel TFs that remodel RSA under combined phosphate and salt stress

We designed a two-stage workflow to identify new transcription factors involved in RSA remodeling under combined low-phosphate and salt stress. The first stage was a bioinformatic analysis that mined public data to surface genes and pathways relevant to both stresses; the second was an experimental program to validate the proposed elements (Figure 1). This two-stage design reflects a deliberate trade-off between breadth and rigor. Bioinformatic mining first cast a wide net across the wealth of already-generated public RNA-seq data, allowing us to survey Pi and salt responses at a scale that no single experiment could achieve, and to statistically prioritize genes and TFs whose regulatory signatures were reproducible across independent projects rather than specific to one dataset. This computational stage, however, can only nominate candidates: co-expression and network centrality indicate association, not causation, and AlphaFold-based interaction modeling predicts physical plausibility. The second, experimental stage was therefore designed to test these predictions directly, using loss-of-function mutants to ask whether the nominated TFs are in fact required for RSA remodeling under combined stress, and whether their phenotypic contributions align with the roles inferred from the network and structural analyses. Together, the two stages form a funnel, from thousands of genes surveyed in silico to a small set of mechanistically validated regulators (Figure 1).

**Figure 1.**
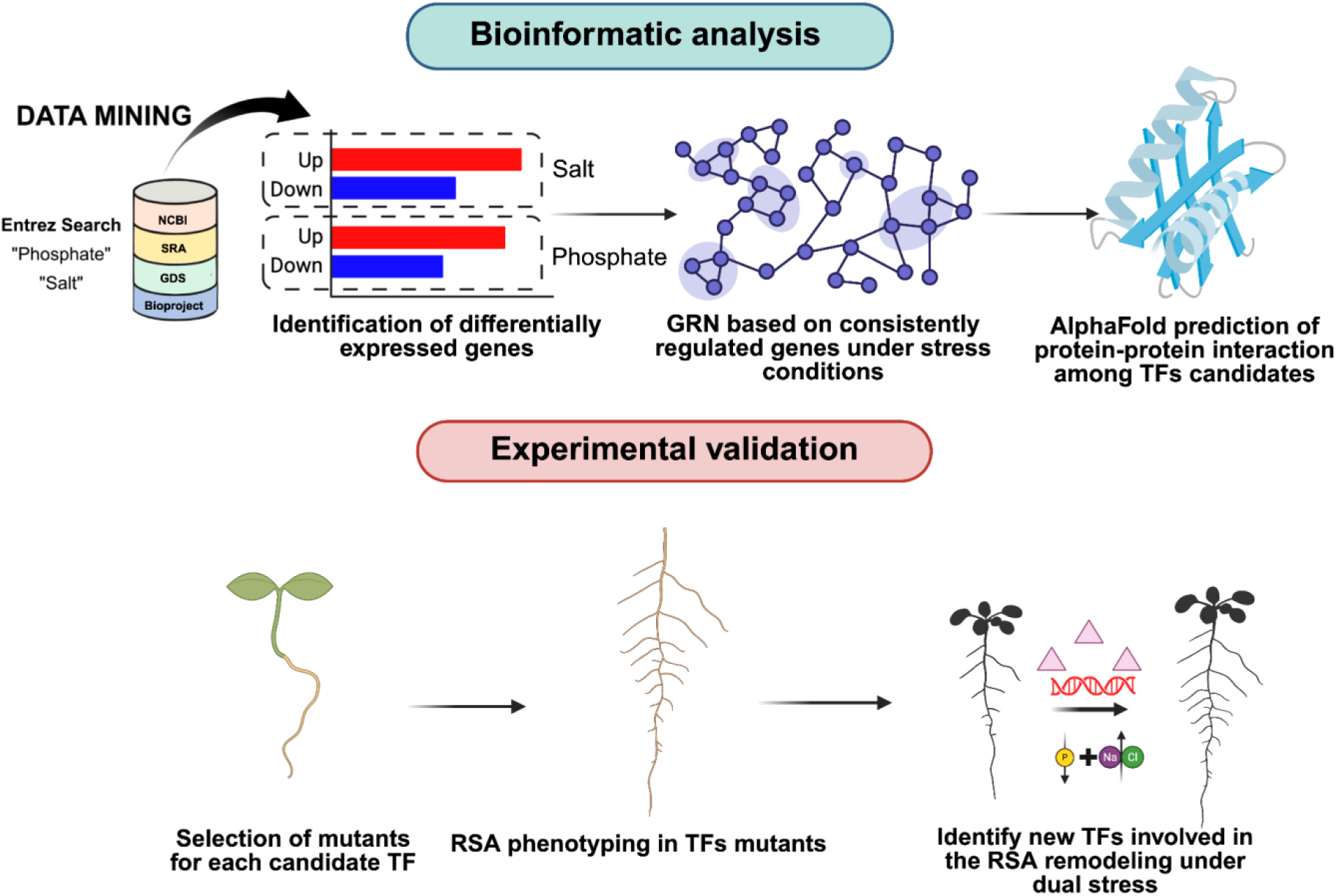
Integrated strategy for identifying transcription factors involved in root system architecture (RSA) remodelling under combined phosphate deficiency and salt stress. *Upper half, bioinformatic analysis.* Public sequencing libraries were retrieved from NCBI by Entrez searches for “phosphate” and “salt” across the BioProject, BioSample, SRA and GDS resources, and differentially expressed genes (DEGs) were called for each control-versus-treatment comparison under each stress. Genes reproducibly regulated across independent comparisons were defined as shared DEGs (sDEGs) and used to build a gene regulatory network (GRN) for each stress. Protein–protein interactions among the top-ranked candidate transcription factors (TFs) of that network were then predicted with AlphaFold-Multimer. *Lower half, experimental validation.* Loss-of-function mutants were selected for each candidate TF, phenotyped for RSA traits under combined low phosphate and salt stress, and used to identify the TFs required for RSA remodelling under the dual stress.

Specifically, we first sought the genes lying at the intersection of the phosphate (Pi) and salt responses in A. thaliana through an integrative analysis of publicly available RNA-seq datasets. Data mining consisted of keyword searches across multiple NCBI repositories (SRA, BioSample, BioProject, GDS), which were merged into complete per-library records. After keyword and manual filtering, we recovered 32 studies (16 each for Pi and salt; Figure 2A), corresponding to 210 (Pi) and 204 (salt) sequencing libraries including treatments and controls. Pi treatments were defined as Pi deprivation—generally growth in trace (≤ 50 µM) or reduced phosphate—and salt treatments as salt addition, generally 75–200 mM NaCl-supplemented medium.

**Figure 2.**
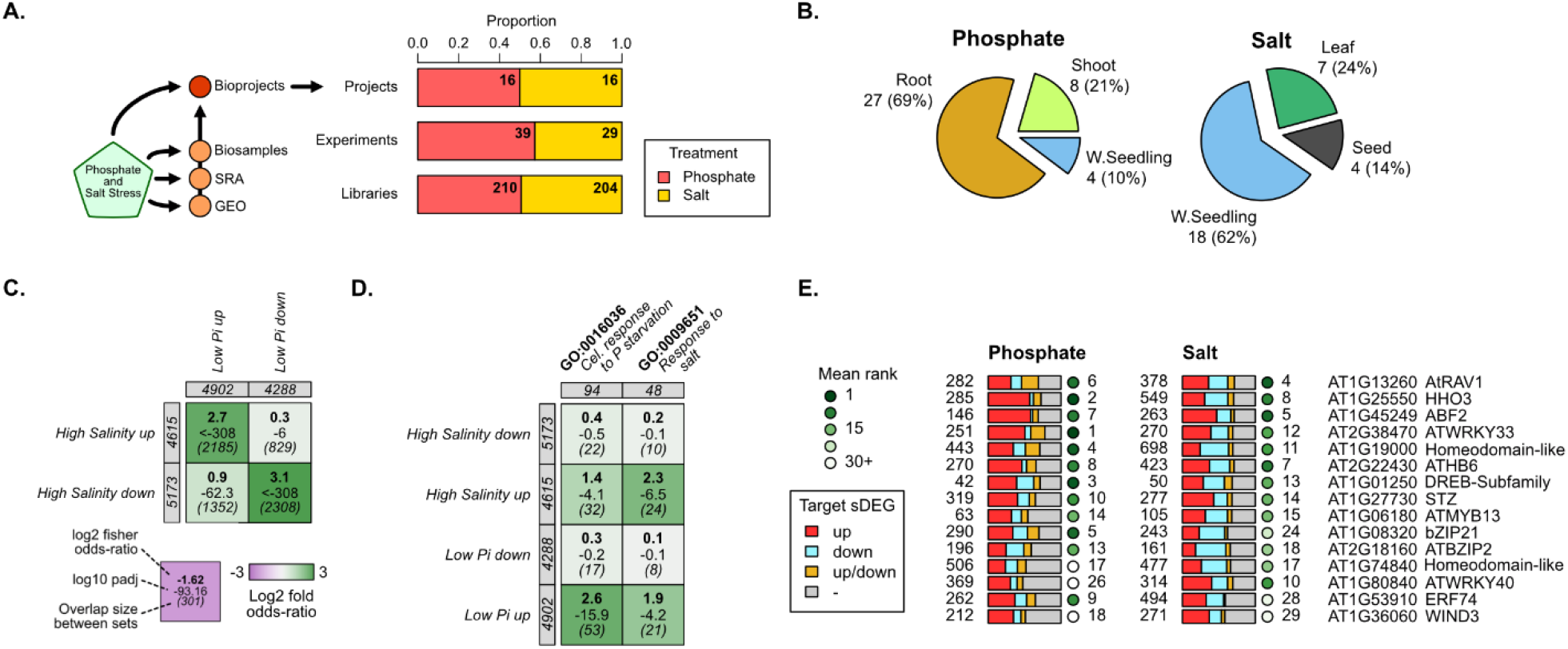
Comparative transcriptomic meta-analysis of phosphate and salt stress identifies candidate transcription factors for RSA remodelling. (A) Data-mining strategy and composition of the compiled dataset. The schematic shows the NCBI resources queried (BioProject, BioSample, SRA and GDS). Stacked bars give the proportion contributed by each treatment at three levels — BioProjects, pairwise control-versus-treatment comparisons (“experiments”) and sequencing libraries; absolute counts are printed inside the bars: 16 phosphate and 16 salt projects, 39 phosphate and 29 salt experiments, and 210 phosphate and 204 salt libraries. (B) Exploded pie charts giving the number of experiments contributed by each tissue, with the percentage of the treatment total in parentheses. The tissue categories differ between treatments: root (27; 69%), shoot (8; 21%) and whole seedling (4; 10%) for phosphate; whole seedling (18; 62%), leaf (7; 24%) and seed (4; 14%) for salt. (C) Gene-set comparison between the high-salt and low-phosphate sDEG sets in all four direction-by-direction combinations, including the two discordant contrasts. Set sizes are printed against the row and column margins (low Pi up, 4,902; low Pi down, 4,288; high salinity up, 4,615; high salinity down, 5,173). Each cell reports three values, decoded by the key below the panel: the log2 fold odds ratio (top, bold), the log10 adjusted P value (middle) and the overlap size between the two sets (bottom, in parentheses). Enrichment was tested by one-sided Fisher’s exact test with Benjamini–Hochberg correction; colour gives the log2 fold odds ratio on the scale at right (−3 to +3, purple to green). (D) As in (C), but comparing each sDEG set against the genes annotated with the Gene Ontology terms “cellular response to phosphate starvation” (GO:0016036; 94 genes) and “response to salt stress” (GO:0009651; 48 genes). Enrichment is markedly stronger for the induced than for the repressed sDEG sets under both stresses. (E) Transcription factors ranked by their out-degree towards sDEGs, shown separately for phosphate and salt. Each horizontal bar is one TF; bar segments give the sDEG status of that TF’s targets — up-regulated (red), down-regulated (light blue), bidirectionally regulated (gold) and not an sDEG (grey). The number at the left of each bar is the out-degree towards sDEGs for that treatment. The filled circle at the right of each bar gives that TF’s rank within that treatment, shaded from rank 1 (filled dark green) through rank 15 to rank 30 or beyond. AGI codes and gene symbols are given at the right. TFs are ordered by the mean of the two per-treatment ranks.

We next examined the relationship between the low Pi and high salinity sDEG sets. Approximately 45–54% of sDEGs were shared between the two stresses (Figure 2C), comprising 2,185 commonly up- regulated and 2,308 commonly down-regulated genes. Set-enrichment analysis (Fisher’s exact test) confirmed a highly significant overlap, displayed as a genesect heatmap (Katari et al., 2010; Figure 2C). Comparing our sDEG sets with the Gene Ontology terms “cellular response to phosphate starvation” (GO:0016036; 94 genes) and “response to salt stress” (GO:0009651; 48 genes) revealed comparable enrichment for each set, with notably stronger enrichment among induced sDEGs (Figure 2D). These results indicate that Pi deficiency and salt stress, despite arising from distinct environmental cues, converge on a substantially overlapping transcriptional program.

Leveraging the breadth of mined libraries, we then built gene regulatory networks for each stress with GENIE3, which benefits from large library counts (> 100 libraries). Raw networks were filtered to retain the top 10% of regulatory relationships per target gene by weight, and then restricted to interactions supported by DAP-seq data (Supplementary Table S1). For each stress, TFs were ranked by their out-degree toward sDEGs (Supplementary Table S2 and S3); the two rankings were then averaged and reordered to surface TFs relevant to both Pi and salt (Figure 2E). The resulting list recovered many established Pi and salt regulators and highlighted candidates that may coordinate crosstalk between the two stresses. Top-ranked genes included well-known phosphate-response regulators such as HHO3 and ABF2 alongside TFs with documented roles in salinity, including DREB- and WRKY-family members.

### AlphaFold protein–protein analysis reveals heterodimerization among ABF2, bZIP2, and HB6

The GRN analysis produced a large list of candidate TFs, which we narrowed to the 15 top-ranked candidates (Figure 2E). Transcription factors are well-established mediators of plant responses to biotic and abiotic stress, frequently acting in concert—often through direct physical interaction—to amplify or repress signaling cascades that help plants endure stress; this has been reviewed by Bhoite et al. (2025). Therefore, to map the protein–protein interaction (PPI) landscape among these candidates, we conducted a systematic in silico structural screen using AlphaFold-Multimer (AFM) (Evans et al., 2022; Kim et al., 2026) (Figure 3). To ensure robust identification of true interacting partners, the confidence and physical extent of the predicted interactions were evaluated using three primary structure-derived metrics: the integrated Local Interaction Score (iLIS), the interface predicted Template Modeling score (ipTM), and the integrated Local Interaction Area (iLIA). iLIS is the geometric mean of the Local Interaction Score (LIS) and its contact-filtered counterpart (cLIS), iLIS = √(LIS × cLIS). LIS is derived from the Predicted Aligned Error (PAE) between residues of opposing chains, averaging the confidence of all residue pairs with PAE ≤ 12 Å; cLIS restricts this to pairs that are additionally in direct physical contact (Cβ–Cβ ≤ 8 Å). A high iLIS indicates that the structural algorithm is highly confident in the relative 3D positioning of the interacting domains. ipTM evaluates the overall topological accuracy of the multimeric assembly at the interface. iLIA is the corresponding integrated interface area, iLIA = √(LIA × cLIA), where LIA counts the confidently predicted residue pairs and cLIA counts the subset in direct contact; it distinguishes transient ’grazing’ contacts from extensive, stable binding surfaces ( Kim et al., 2026).

**Figure 3.**
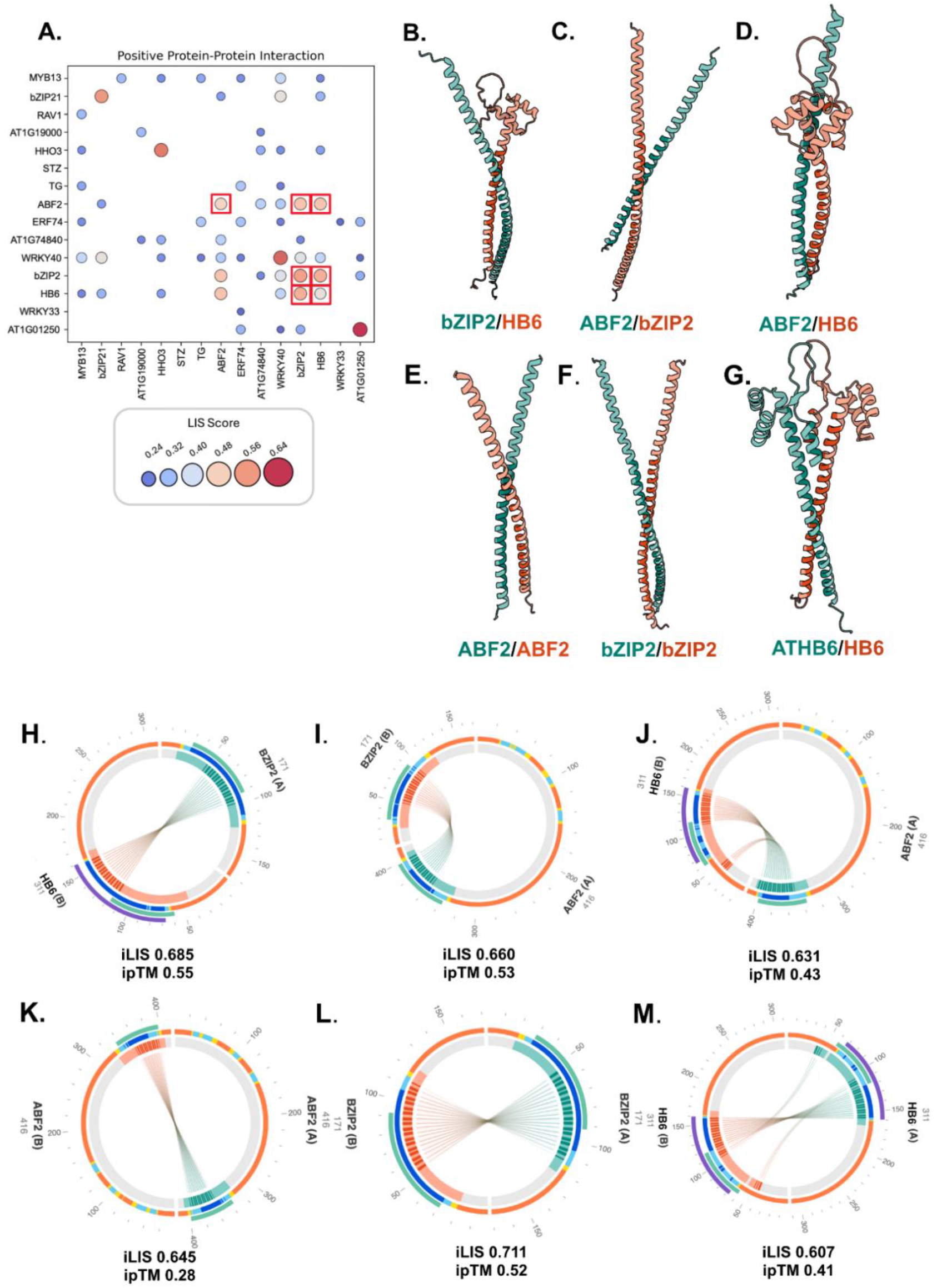
AlphaFold-Multimer structural screening reveals an interconnected protein–protein interaction network and a distinct ABF2–bZIP2–HB6 module among the top candidate transcription factors. (A) Pairwise interaction matrix for the 15 top-ranked candidate transcription factors of Figure 2E. All 120 possible complexes were modelled (105 heterotypic pairs plus 15 homodimers); only pairs classified as positive interactions at the threshold given in the Methods are plotted. Bubble area and colour give the best score across five independent models, on the scale shown below the matrix (0.24– 0.72). Homotypic interactions lie on the diagonal; off-diagonal bubbles are predicted heterodimers. Red boxes mark the interactions within the ABF2–bZIP2–HB6 sub-matrix. (B–G) Predicted three-dimensional structures of the highest-ranked model for each complex: bZIP2/HB6 (B), ABF2/bZIP2 (C), ABF2/HB6 (D), ABF2/ABF2 (E), bZIP2/bZIP2 (F) and HB6/HB6 (G). The two chains are shown in teal and salmon. All six complexes are predicted to form intertwined helical coiled-coil interfaces. Structures were rendered in ChimeraX (Meng et al., 2023). (H–M) Predicted residue-level interaction interfaces for the same six complexes, in the same order: bZIP2/HB6 (H), ABF2/bZIP2 (I), ABF2/HB6 (J), ABF2/ABF2 (K), bZIP2/bZIP2 (L) and HB6/HB6 (M). In each circular plot the two chains are laid out end to end around the circumference, labelled with the protein name, the chain identifier (A or B) and the residue number at the chain terminus (ABF2, 416 residues; bZIP2, 171; HB6, 311). The outer arcs represent the individual chains. The inner concentric tracks display the per-residue prediction confidence (pLDDT) mapped to a colour scale: dark blue (very high, >90), light blue (high, 70–90), yellow (low, 50–70), and orange (very low, <50). Central chords connect cLIR (contact Local Interaction Residue) pairs predicted to be in physical contact at the interface. The integrated Local Interaction Score (iLIS) and interface predicted Template Modelling score (ipTM) of the best model are given below each plot: bZIP2/HB6, iLIS 0.685, ipTM 0.55; ABF2/bZIP2, iLIS 0.660, ipTM 0.53; ABF2/HB6, iLIS 0.631, ipTM 0.43; ABF2/ABF2, iLIS 0.645, ipTM 0.28; bZIP2/bZIP2, iLIS 0.711, ipTM 0.52; HB6/HB6, iLIS 0.607, ipTM 0.41. Both homodimeric and heterodimeric configurations of ABF2 and bZIP2 are therefore predicted with comparable confidence, the bZIP2 homodimer scoring highest of the six.

For each protein pair, AlphaFold-Multimer generated five independent model simulations. Interaction parameters were calculated for the best-scoring model ("Best") and as a mean across all five models ("Average"). Evaluating the "Average" scores provides a measure of structural reproducibility, confirming that the complex folds consistently across different computational runs, whereas the "Best" score captures the most structurally favorable conformation achieved. Requiring high values for these interface-specific metrics, rather than relying on global AlphaFold-Multimer scores alone, provides a stricter and more reliable threshold for distinguishing genuine interactors among the 15 screened TF pairs.

By plotting the ’Best’ and ’Average’ interaction metrics, we established stringent empirical thresholds. While the baseline for a positive interaction was defined by the reference-set-derived threshold of iLIS ≥ 0.223 (Kim et al., 2026), we applied a more stringent, internally calibrated cut-off (iLIS best ≥ 0.551 and iLIS avg ≥ 0.303) to filter for the most robust assemblies. Structural models surpassing this stringent cut- off were classified as highly confident positive PPIs, representing approximately 30% of the 120 complexes modelled (105 heterotypic pairs plus 15 homodimers; Supplementary Table S4). Ranking the pairs by their best iLIS score, the resulting interaction matrix revealed a highly interconnected network comprising both homodimeric and heterodimeric associations (Figure 3A).

While several candidates exhibited a strong propensity to form homodimers (e.g., HHO3, ABF2, bZIP2, HB6, WRKY40, and AT1G01250), a distinct cross-family interaction pattern emerged. Strikingly, only three candidates robustly interacted with one another in a heterotypic manner: ABF2 with bZIP2 (iLIS = 0.660, ipTM = 0.53), ABF2 with HB6 (iLIS = 0.631, ipTM = 0.43), and bZIP2 with HB6 (iLIS = 0.685, ipTM = 0.55).

The corresponding homodimers scored comparably or higher — ABF2/ABF2 (iLIS = 0.645, ipTM = 0.28), bZIP2/bZIP2 (iLIS = 0.711, ipTM = 0.52) and HB6/HB6 (iLIS = 0.607, ipTM = 0.41; Figure 3K–M) — the bZIP2 homodimer being the highest-scoring model in the screen. Both homodimeric and heterodimeric configurations are therefore predicted with high confidence, which is the arrangement required by the dimer-specific target partitioning described below. This indicates that these three TFs may form a mutually interacting, combinatorial regulatory module that coordinates the transcriptional response to combined stress. Structural modeling of each pair further resolved these interactions, demonstrating that the predicted HB6-bZIP2, ABF2-bZIP2, and ABF2-HB6 heterodimers form tightly intertwined helical coiled-coil interfaces with extensive contacts (Figure 3B–D). Similar structural conformations were observed for their respective homodimers (Figure 3E-G). These structural predictions are consistent with a specific physical association, which remains to be tested experimentally.

### Transcriptional regulatory network and dimer-specific target regulation by ABF2 and bZIP2

Having identified ABF2, HB6, and bZIP2 as core, physically interacting components of this stress module, we next asked how they translate this interaction into transcriptional control of the combined Pi and salt response. To map their direct regulatory architecture, we constructed a gene regulatory network (Figure 4A) strictly filtered using in vitro DNA-binding evidence from DNA Affinity Purification sequencing (DAP-seq) assays. The bipartite network topology distinctly partitions the target genes into large, independent ABF2 and bZIP2 regulons on the periphery, anchored by a central core of highly shared targets. By overlaying transcriptomic metadata, we found that a substantial fraction of the network is root-expressed and responsive to low Pi and/or salt stress: of the 277 genes in the network, 105 (38%) are expressed in root or root structures (Figure 4A; Supplementary Table S5). This pervasive stress-responsiveness across both unique and overlapping targets firmly positions ABF2 and bZIP2 as central executioners of the multi-stress root response.

**Figure 4.**
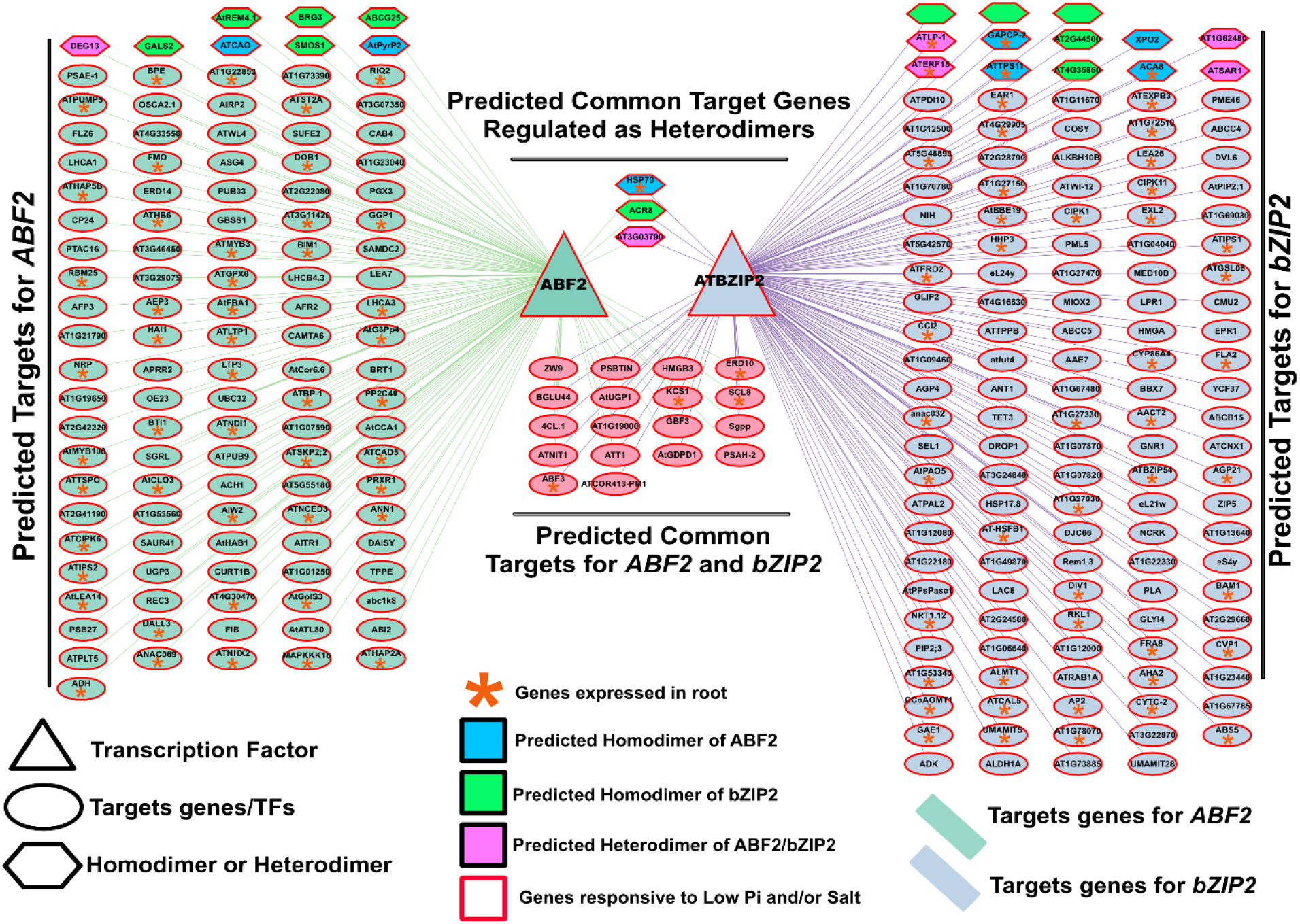
DAP-seq-filtered gene regulatory network of ABF2 and bZIP2 and a dimer-specific model of target regulation. (A) Gene regulatory network of the predicted target genes of ABF2 and bZIP2. Networks were inferred with GENIE3 from the compiled phosphate- and salt-responsive RNA-seq libraries, filtered to the top 10% of interactions per TF by weight, and retained only where supported by DAP-seq binding evidence from the Plant Cistrome Database (O’Malley et al., 2016). Node shape denotes class: triangles, transcription factors; ellipses, target genes; hexagons, targets assigned to a specific dimeric configuration. Node fill denotes the predicted dimer responsible for regulation: blue, ABF2 homodimer; green, bZIP2 homodimer; magenta, ABF2/bZIP2 heterodimer; unfilled, not assigned. A red node outline marks genes responsive to low Pi and/or salt in the meta-analysis, and an orange asterisk marks genes expressed in root or root structures. Edge colour denotes the regulator: green, ABF2 targets; purple, bZIP2 targets. Targets unique to each TF are arrayed at the left (ABF2) and right (bZIP2); the two central groups are the targets shared by both TFs (lower) and the subset predicted to be regulated specifically as heterodimers (upper). Of the 277 genes in the network, 105 (38%) are expressed in root or root structures. Full node annotation is given in Table S5.

Because basic leucine zipper (bZIP) transcription factors canonically bind to DNA as dimers, we sought to determine whether these downstream targets are regulated by homodimeric or heterodimeric configurations (Jakoby et al., 2002). To computationally infer functional dimerization on the DNA, we performed a structural promoter analysis. Specifically, we searched for target promoters containing adjacent binding motifs separated by a constrained gap of 3 to 10 base pairs—an optimal spatial arrangement required to facilitate the simultaneous physical anchoring of bZIP dimers (Schütze et al., 2008; Dröge-Laser et al., 2018). Based on this analysis, we excluded HB6 from the central regulatory node of the network; although it physically interacts with the other factors, the motif-spacing data revealed that HB6 homo- and heterodimers regulate very few target genes, positioning it as a functional outgroup relative to the dense ABF2/bZIP2 regulatory core (Supplementary Table S6). In contrast, this spatial motif analysis allowed us to distinctly classify targets across the core network as being under the homodimeric or heterodimeric control of ABF2 and bZIP2 (Figure 4A).

Integrating this motif-spacing spatial data with our filtered network, we propose a comprehensive regulatory model illustrating the functional interplay between ABF2 and bZIP2 (Figure 4B). First, the model identifies a hierarchical regulatory relationship wherein ABF2 acts directly upstream of bZIP2. More importantly, it highlights a complex, dimer-specific regulatory mechanism for their downstream effectors. Our data robustly demonstrate that distinct dimeric configurations preferentially orchestrate specific targets: AT3G03790 is uniquely regulated by the ABF2 homodimer, ACR8 (AT1G12420) is governed by the bZIP2 homodimer, and HSP70 (AT4G16660) is under the strict combinatorial control of the ABF2/bZIP2 heterodimer. Together, these findings indicate that ABF2 and bZIP2 coordinate stress-responsive gene expression not only through hierarchical control but also via fine-tuned, motif-dependent dimeric partitioning on target promoters.

### Single and double mutants display distinct yet overlapping constraints on root system architecture under combined stress

Single mutants display distinct yet overlapping RSA defects. Macroscopic evaluation of seedlings (Figure 5A) revealed visible root growth defects across the mutant lines. Quantitative analysis via Log₂ fold-change (FC) relative to Col-0 (Figure 5C, top panel) demonstrated that under control conditions, the bzip2 mutant exhibits a strong, constitutive impairment across nearly all RSA traits, with highly significant reductions in primary root length, lateral root length and lateral root number, alongside a significant increase in lateral root density (p < 0.001). This indicates that while *bzip2* initiates fewer lateral roots, they are proportionally denser along the shortened primary root. In contrast, the *abf2* mutant maintained primary and lateral root lengths comparable to the wild-type under control conditions, but displayed a severe defect in lateral root initiation, reflected in highly significant reductions in both lateral root number and density (p < 0.001). Upon exposure to combined stress, *bzip2* showed a relative recovery in primary root length (ns relative to Col-0), while retaining strong lateral root defects. Notably, under the same stress conditions, the *abf2* mutant uniquely displayed a significant increase in lateral root length compared to Col-0 (p < 0.001, Figure 5C), suggesting a compensatory elongation mechanism despite its severely diminished lateral root density.

**Figure 5.**
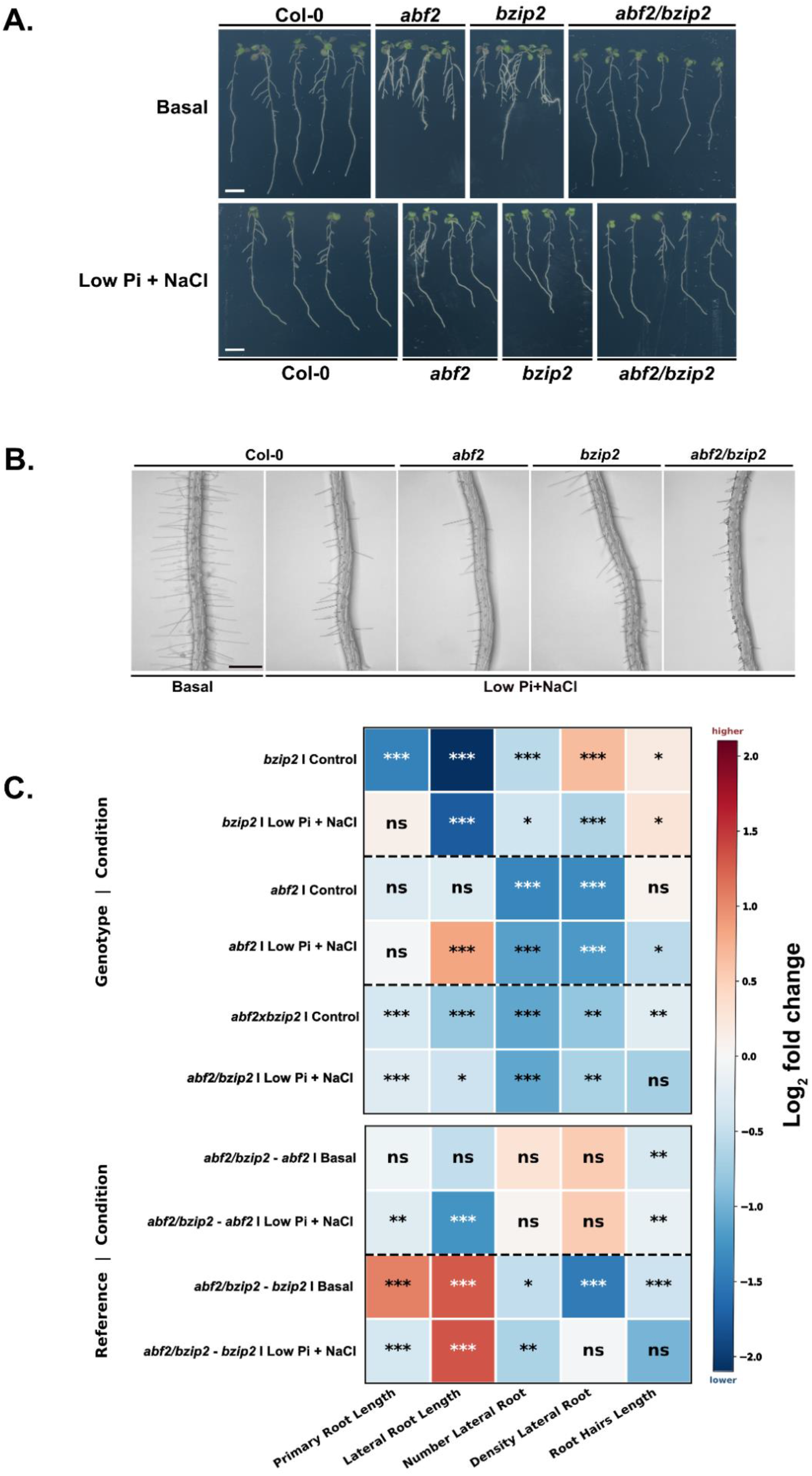
Root system architecture of *abf2*, *bzip2* and *abf2/bzip2* mutants under combined low-phosphate and salt stress. (A) Representative whole-seedling images of Arabidopsis thaliana Col-0, *abf2*, *bzip2* and *abf2/bzip2* grown for 7 d on basal medium (half-strength MS) and then transferred for 3 d to basal medium (Basal, upper row) or to medium containing 5 µM Pi plus 100 mM NaCl (Low Pi + NaCl, lower row); seedlings were therefore ten days old at imaging. Images were acquired on an EPSON Perfection V8 scanner. Scale bars = 0.2 cm. (B) Representative micrographs of the elongation/differentiation zone of the primary root of the same four genotypes, showing root hair development under basal conditions (Col-0, left) and under combined low Pi + NaCl (all four genotypes, right). Images were acquired on a Leica EZ4 HD stereomicroscope with LAS EZ software. Scale bar = 600 µm. (C) Heatmap of Log2 fold changes in five RSA traits — primary root length, lateral root length, lateral root number, lateral root density and root hair length (columns, left to right). Upper block (Genotype | Condition): each mutant relative to Col-0 within the same treatment. Lower block (Reference | Condition): the *abf2/bzip2* double mutant relative to each single mutant within the same treatment. Red indicates a higher and blue a lower trait value than the reference, on the scale at right (−2.0 to +2.0). Each genotype was compared with its own contemporaneous Col-0 control, since Col-0 absolute values differ between independent experiments (see Fig. S4). Data are the mean of 30 seedlings per genotype per condition (10 seedlings × 3 independent biological replicates); root hair values are the longest 10 root hairs from each seedling, this means that are 10 root hairs per plant. Significance was assessed by two-way ANOVA followed by Tukey’s multiple-comparison test in GraphPad Prism v8.0.1. Asterisks denote the difference from the stated reference within each treatment: *P < 0.05; **P < 0.01; ***P < 0.001; ****P < 0.0001; ns, not significant. Source data are shown per trait in Figs S4–S9.

Simultaneous loss of both transcription factors resulted in a robust and constitutive repression of the root system. As observed visually in the whole-seedling macro-phenotype (Figure 5A) and quantitatively in the fold-change heatmap (Figure 5C), the *abf2/bzip2* double mutant displayed highly significant negative fold-changes across all measured macroscopic RSA traits under control conditions. Specifically, the double mutant showed a 22.8% reduction in primary root length (2.78 vs. 3.60 cm in Col-0; p < 0.0001) and a drastic 53% reduction in lateral root number (3.3 vs. 7.0 roots; p < 0.0001). When normalized to primary root length, lateral root density remained significantly compromised (1.14 vs. 2.00 roots/cm; p < 0.01), indicating a fundamental defect in lateral root emergence rather than a secondary consequence of reduced primary root elongation. Under combined Low Pi + NaCl stress, this systemic inhibition persisted: the double mutant maintained a significantly shorter primary root (17% reduction; p < 0.001) and a severely diminished lateral root branching network (53% reduction in number; p < 0.001) relative to stressed wild-type plants. Furthermore, comparative Log₂ fold-change analysis between the double mutant and the respective single mutants (Figure 5C, bottom panel) highlights the synergistic nature of these defects, as the double mutant displays significantly exacerbated reductions in lateral root formation and density compared to either the *abf2* or *bzip2* mutation alone.

Microscopic examination (Figure 5B) revealed that while Col-0 produces long and dense root hairs under control conditions, combined Low Pi + NaCl treatment restricts root hair development. Under basal conditions, the double mutant showed a moderate, yet significant, reduction in root hair length relative to absolute Col-0 levels (312 µm vs. 371 µm; 16% reduction; p < 0.01). Under combined stress, this absolute impairment became severe: *abf2/bzip2* root hairs averaged merely ∼57 µm, representing a drastic and statistically significant 47.3% reduction in absolute length relative to the ∼108 µm observed in stressed Col-0 (p < 0.001; Supplementary Figure S9E). However, when evaluated as a stress-induced fold-change relative to their respective basal controls, the proportional reduction in the double mutant did not reach statistical significance against the wild-type response (ns, p = 0.0662; Figure 5C heatmap). Taken together, these results visually and quantitatively demonstrate that ABF2 and bZIP2 act in an additive or synergistic manner to sustain not only macroscopic root branching but also the critical absolute elongation of root hairs required for acclimation to combined nutrient deprivation and osmotic stress.

## Discussion

Climate change increasingly compromises crop productivity, exacerbating the challenge of plant adaptation to degraded and saline soils. Understanding how plants integrate multiple, simultaneous stresses—rather than individual stresses in isolation—is essential. Although responses to phosphate limitation and salinity have each been studied extensively, their combination has received far less attention despite routinely co-occurring in the field. To address this, we deployed an integrative pipeline: public-data meta-analysis, gene regulatory network (GRN) inference, structure-based interaction prediction, and mutant phenotyping. This approach offers a generalizable template for dissecting other combined-stress scenarios.

A central observation from our meta-analysis is the extensive overlap between the phosphate and salt transcriptional programs. Roughly 45–54% of the shared differentially expressed genes (sDEGs) were common to both stresses, and this overlap was statistically significant for both induced and repressed genes. This convergence argues that the two stress responses are not independent but instead share a substantial regulatory architecture, consistent with prior reports of crosstalk between nutrient and ionic-stress signaling (Khan et al., 2023). By retaining only those transcription factor (TF)-target interactions supported by DAP-seq binding evidence, our networks successfully recovered numerous canonical phosphate and salinity regulators. This recovery of known players provides internal validation and lends high confidence to the novel candidates that emerged alongside them.

While interactions between TF families are well documented—particularly the importance of homo- and heterodimerization in fine-tuning stress responses (Schütze et al., 2008; Dröge-Laser et al., 2018)—there is limited structural and network-level evidence regarding how these complexes regulate diverse gene sets under combined stress. Our AlphaFold-Multimer analysis predicted that ABF2, bZIP2, and HB6 form a mutually interacting module. Biologically, this is highly coherent: ABF2 and bZIP2 are basic leucine-zipper (bZIP) factors central to abscisic acid (ABA) signaling, while HB6 is a homeodomain-leucine-zipper (HD-Zip) factor and a documented modulator of ABA responses (Söderman et al., 1999; Himmelbach et al., 2002). Because ABA is a key hormonal node linking osmotic and ionic stress to phosphate-status signaling, a physical hub bringing together bZIP and HD-Zip regulators offers an attractive mechanism for coordinated stress integration. However, when we transitioned from protein-protein interaction predictions to hierarchical GRN construction, the functional roles of these TFs on the DNA diverged. Using DAP-seq data combined with a stringent motif-spacing filter—requiring adjacent binding motifs separated by 3 to 10 base pairs to accommodate dimer anchoring—we discovered that HB6 acts as a regulatory outgroup. Despite structurally interacting with the module, HB6 dimers directly regulate very few targets within the shared stress network. Consequently, our focus narrowed to the highly connected ABF2 and bZIP2 core. Within this central node, our GRN hierarchically positioned ABF2 upstream, directly regulating bZIP2. Furthermore, filtering for targets governed strictly by dual motifs highlighted key downstream effectors under specific homo- or heterodimeric control. For instance, the central node includes *ACR8* (AT1G12420), a target with a previously uncharacterized biological role; *HSP70* (AT4G16660), a cAMP-responsive gene involved in ABA signaling during dehydration (Babar et al., 2026); and AT3G03790, which root atlas tools reveal is highly expressed in the quiescent center, procambium, and, critically, trichoblasts and atrichoblasts.

Crucially, the identification of a trichoblast-expressed target provides a direct molecular link to our phenotypic data, as trichoblasts are the epidermal cells responsible for root hair formation. Furthermore, the reliance of specific targets like *HSP70* (AT4G16660) strictly on the ABF2/bZIP2 heterodimer provides a mechanistic basis for the synergistic morphological paralysis observed in our double mutant. While single mutants may retain partial functionality through their respective homodimers, the double mutant suffers a complete collapse of both homo- and heterodimeric regulatory arms. This strong root-specific expression profile is mirrored across the broader network: of the 277 genes composing the core GRN, 105 (38%) are expressed in root or root structures. The compiled dataset is itself root-weighted: root material accounted for 69% of the phosphate experiments and whole seedlings for 62% of the salt experiments (Figure 2B). This transcriptomic bias strongly suggested that the root system is the primary morphological arena for this combined stress response, prompting our phenotypic analyses.

Our mutant analyses robustly validated the regulatory roles of these factors, revealing that ABF2 and bZIP2 shape distinct yet overlapping subsets of RSA. Biologically, these two stresses impose conflicting demands on root architecture: phosphate starvation typically stimulates lateral root and root hair proliferation to maximize soil foraging, whereas salinity generally represses these traits to prevent the uptake of toxic ions. The near-complete arrest of root hair elongation and lateral root formation in the *abf2/bzip2* double mutant under combined stress positions this bZIP module as a critical regulatory "switch" required to balance these conflicting signals. By sustaining specific dimensions of root growth, these factors allow for essential nutrient acquisition despite osmotic stress. That these single-gene knockouts, and particularly the double mutant, produce such measurable and severe phenotypes—despite probable genetic redundancy among the broader bZIP family—underscores their indispensable functional importance in multi-stress acclimation.

Several limitations temper these conclusions. First, meta-analysis of heterogeneous public datasets inevitably aggregates variation in genetic background, growth systems, stress severity, and sampling times. While our stringent requirements for cross-experiment consistency were designed to mitigate this variability, it cannot be fully eliminated. Second, our phenotypic assays were conducted in vitro on agar plates with a defined combined-stress regime imposed after seedling establishment. While these conditions accurately capture early adaptive responses, they only partially reflect the chronic, fluctuating stresses of field soils. Third, the RSA phenotyping compared basal medium with the combined stress only; low-Pi and NaCl single-stress arms were not phenotyped, so the extent to which the combined response differs from the salt response alone cannot be resolved here. Finally, our AlphaFold models identify highly plausible structural interactions rather than experimentally proven in vivo complexes. Future work integrating in planta interaction assays (such as bimolecular fluorescence complementation), higher-order mutants to address broader family redundancy, direct target validation by RT-qPCR and chromatin profiling, and phenotypic assessment under soil-based combined stress will be essential. Ultimately, this will convert these candidate networks into a validated mechanistic model of combined-stress adaptation with direct translational value for breeding climate-resilient crops.

## Conclusions

By integrating a meta-analysis of publicly available transcriptomic data with gene regulatory network inference, AlphaFold-based interaction prediction, and functional genetics, we identified ABF2, bZIP2 and HB6 as transcription factors predicted to interact physically during root system architecture remodeling under combined low-phosphate and salt stress. The pronounced overlap between the phosphate and salt transcriptional programs indicates extensive crosstalk between these stresses. While structural modeling highlights an ABA-anchored physical interaction network among these three factors, our DNA-motif and phenotypic analyses established the ABF2–bZIP2 dimer pair as the core regulatory hub through which these dual stress inputs are integrated. Loss-of-function mutants of ABF2 and bZIP2, singly and in combination, displayed severely altered RSA, with each gene shaping a distinct set of traits (characterization of the *hb6* single mutant is provided in Supplementary Figures S4–S8). Beyond the specific regulators uncovered here, this study demonstrates the value of combining data mining, structural prediction, and targeted phenotyping to dissect multifactorial stress, and it lays the groundwork for engineering root traits that improve crop resilience on the increasingly phosphate-poor and saline soils of a changing climate.

## Supplementary data

The following supplementary data are available at *JXB* online.

**Fig. S1**. Definition of shared differentially expressed genes (sDEGs) across the compiled phosphate and salt experiments.

**Fig. S2.** Overview of the phosphate and salt treatment conditions represented in the compiled dataset.

**Fig. S3.** Distribution of DEGs across individual experiments and transcription factor content of the sDEG classes.

**Fig. S4.** AlphaFold-Multimer prediction of protein–protein interactions among ABF2, bZIP2, and ATHB6.

**Fig. S5**. Gene expression of each TF (*ABF2*, *HB6,* and *bZIP2*) in the main root and in the root cell type.

**Fig. S6.** Primary root length of *abf2*, *bzip2*, *hb6 and double mutant (abf2/bzip2)* under combined low phosphate and salt stress.

**Fig. S7.** Lateral root length of *abf2*, *bzip2*, *hb6 and double mutant (abf2/bzip2)* under combined low phosphate and salt stress.

**Fig. S8**. Number of lateral roots of *abf2*, *bzip2*, *hb6 and double mutant (abf2/bzip2)* under combined low phosphate and salt stress.

**Fig. S9**. Lateral root density of *aabf2*, *bzip2*, *hb6 and double mutant (abf2/bzip2)* under combined low phosphate and salt stress.

**Fig. S10.** Root hair length of *aabf2*, *bzip2*, *hb6 and double mutant (abf2/bzip2)* under combined low phosphate and salt stress.

**Fig. S11.** Working model: ABF2–bZIP2 transcriptional cooperation mediates root architecture remodelling under combined low phosphate and salt stress.

**Table S1**. Sequencing libraries, BioProject accessions and metadata retained after filtering.

**Table S2.** Transcription factors ranked by out-degree towards sDEGs for Phosphate.

**Table S3.** Transcription factors ranked by out-degree towards sDEGs for Salt.

**Table S4**. AlphaFold-Multimer structural screening scores. Description: Integrated Local Interaction Scores (iLIS) and interface predicted Template Modeling (ipTM) scores for all 120 modelled transcription factor pairs.

**Table S5**. List of the 277 genes in the low Pi–salt network, categorized according to their root expression status.

**Table S6**. Selection of heterodimeric and homodimeric motifs for ABF2, bZIP2, and HB6.

## Acknowledgements

We thank ABRC for seed stocks, the Harvard O2 high-performance computing cluster for the AlphaFold-Multimer runs.

## Author contributions

JMA and JME: conceptualization; NRJ and TCM: methodology, software and formal analysis; HS-G and MAI: investigation; JP-D, RA-E, LA, VBG and GV-M: investigation and validation; NRJ and TCM: data curation; JMA and JME: writing – original draft; all authors: writing – review and editing; JMA and JME: supervision; JMA and JME: funding acquisition. All authors read and approved the final version.

## Data availability

All transcriptomic data analysed in this study are publicly available from the NCBI Sequence Read Archive; the BioProject and run accessions for the 32 studies retained after filtering are listed in Supplementary Table S1. DAP-seq data were obtained from the Plant Cistrome Database (O’Malley et al., 2016). Analysis code is available and the AFM-LIS implementation used here is at github.com/flyark/AFM-LIS and the genesectr package at github.com/nateyjay/genesectr. All other data supporting the findings are available from the corresponding authors on reasonable request.

## Abbreviations

DAP-seq: DNA affinity purification sequencing
DEG: differentially expressed gene
GRN: gene regulatory network
iLIS: integrated Local Interaction Score
LR: lateral root
PR: primary root
RH: root hair
RSA: root system architecture
sDEG: shared differentially expressed gene
TF: transcription factor.

